# Nicotine self-administration increases impulsive action: differential effects of nAChR modulators in a Go/No-Go task

**DOI:** 10.64898/2026.03.31.715632

**Authors:** Ranjithkumar Chellian, Guido Huisman, Adriaan W. Bruijnzeel

## Abstract

Tobacco use disorder is a chronic condition characterized by compulsive nicotine use, withdrawal, and relapse following abstinence. Impulsivity contributes to persistent nicotine use and poor cessation outcomes. This study examined whether nicotinic acetylcholine receptor (nAChR) modulators alter impulsive action in a nicotine self-administration Go/No-Go task in male and female rats. Rats acquired intravenous nicotine self-administration and were then trained in a Go/No-Go procedure in which active lever presses were reinforced during Go periods but not during No-Go periods. We then assessed the effects of varenicline (0.1-3 mg/kg), nicotine (0.1-0.6 mg/kg), and the nAChR antagonist mecamylamine (0.5-2 mg/kg) in the Go/No-Go procedure. Varenicline and nicotine pretreatment reduced active responding during both Go and No-Go periods, whereas mecamylamine selectively reduced responding during No-Go periods. Mecamylamine decreased the percentage of active responses during No-Go trials, indicating reduced bias toward the nicotine-associated lever. In contrast, nicotine and varenicline did not alter response allocation, suggesting that their effects reflected nonspecific reductions in responding rather than changes in impulsive action. No sex differences were observed. Substituting saline for nicotine during self-administration did not alter active responding during Go periods, but rats in the saline group had fewer active responses during No-Go periods than rats in the nicotine group. These results show that chronic nicotine self-administration increases impulsive action and that nAChR antagonism, but not agonism or partial agonism, reduces nicotine-related impulsive action. This work supports the utility of the Go/No-Go self-administration task for investigating nAChR-dependent mechanisms underlying nicotine-induced impulsivity.

## 1. Introduction

Tobacco use disorder is a chronic condition characterized by the persistent, problematic use of tobacco products that disrupts daily functioning and leads to emotional distress (American Psychiatric Association 2013). Specifically, smokers may experience a loss of control over tobacco use, a persistent desire or unsuccessful efforts to quit, craving, continued use despite adverse consequences, and withdrawal (American Psychiatric Association 2013). Worldwide, approximately 1.25 billion people smoke, and tobacco use remains the leading preventable cause of morbidity and mortality (WHO 2024). Smoking is associated with a broad spectrum of diseases, including cardiovascular disease, respiratory disease, cancer, and Alzheimer’s disease (Ambrose and Barua 2004; Forey et al. 2011; Ott et al. 1998; Sasco et al. 2004). In addition to its adverse health consequences, tobacco use is a significant economic burden due to an increase in medical expenses and lost productivity (Goodchild et al. 2018; Nargis et al. 2022). Despite the availability of FDA-approved smoking cessation aids and widespread public health initiatives, smoking and relapse rates remain high (Agboola et al. 2015; Cornelius et al. 2023; Williams et al. 2007).

Impulsivity plays a critical role in the initiation, maintenance, and relapse of tobacco use (Doran et al. 2004; Perry and Carroll 2008). Impulsivity is a tendency toward rapid, often poorly considered actions that can have negative consequences for oneself or others (Moeller et al. 2001). Impulsivity encompasses several distinct components, including response inhibition, delay discounting, and attentional impulsivity (Jauregi et al. 2018; Weafer et al. 2011). A deficit in the inhibition of responses has been associated with substance abuse, including tobacco use disorder (Smith et al. 2014). The Go/No-Go task has been widely used in humans and animal studies to assess inhibitory control (Casey et al. 1997; Durston et al. 2002; Gubner et al. 2010; Rubia et al. 2001), specifically by measuring the inhibition of motor responses before they begin (Eagle et al. 2008). Tobacco users have diminished inhibitory control in the Go/No-Go task compared to nonsmokers (Luijten et al. 2011). In Go/No-Go procedures, responding during No-Go periods reflects failures of response inhibition (impulsive action). Additionally, the percentage of active responses is commonly used as an index of response bias in impulsivity and behavioral inhibition studies (Eagle et al. 2008; Kolokotroni et al. 2011; Winstanley 2011). Pharmacological treatments, neuromodulation techniques, and behavioral therapies that improve impulse control have shown efficacy in reducing substance use and relapse among individuals with substance use disorders (Goldstein and Volkow 2011; Steele et al. 2019; Zilverstand et al. 2018). For instance, transcranial direct current stimulation (tDCS) has shown promise in enhancing inhibitory control and decreasing relapse rates in people with alcoholism by modulating activity in the dorsolateral prefrontal cortex (Dubuson et al. 2021).

Nicotine is a nonselective agonist at nicotinic acetylcholine receptors (nAChRs) and mediates its rewarding effects by activating α4β2, α3β4, α7, and α6-containing nAChRs (Picciotto and Kenny 2021). Animal models play a critical role in improving our understanding of the neurobiological mechanisms underlying nicotine self-administration, withdrawal, and relapse (Chellian et al. 2021a). Rats readily learn to self-administer nicotine, and nicotine has rewarding properties in well-established assays such as conditioned place preference and intracranial self-stimulation (Chellian et al. 2025; Chellian et al. 2021b; Corrigall and Coen 1989; Le Foll and Goldberg 2005). Furthermore, cessation of nicotine intake leads to withdrawal and animals display nicotine seeking behavior after a period of abstinence (Chellian et al. 2024b). Animal models have also been developed to study impulsivity and response inhibition and allow for the investigation of the neurobiological mechanisms that mediate increased impulsivity in smokers (Eagle et al. 2008; Winstanley 2011). The rodent Go/No-Go task requires animals to discriminate between cues signaling reward delivery (Go) and cues indicating reward omission (No-Go) and thereby enables the assessment of inhibitory control. Kolokotroni et al. assessed impulsivity using a symmetrically reinforced Go/No-Go task, in which both correct responses during Go trials and successful inhibition during No-Go trials were rewarded with sucrose pellets (Kolokotroni et al. 2011). They found that acute noncontingent nicotine administration impaired response inhibition in rats (Kolokotroni et al. 2011).

Our study expands upon previous research by investigating the effects of chronic nicotine self-administration in both male and female Wistar rats, offering a more translatable model for tobacco use disorders than prior studies using acute nicotine administration to drug-naïve animals (Kolokotroni et al. 2011). In our study, we adapted a Go/No-Go procedure previously used in cocaine self-administration research to examine the effects of nicotine self-administration on impulsive action (Zapata et al. 2017; Zapata and Lupica 2021). Rats were trained to self-administer nicotine, and we assessed nicotine intake during the Go periods and lever responding during the No-Go periods as a measure of impulsive action. To investigate the role of FDA-approved smoking cessation treatments and nAChRs in impulsive behavior, we examined the effects of pretreatment with varenicline (Chantix), mecamylamine, and nicotine on performance in the Go/No-Go task. Varenicline is a partial agonist at α4β2 nAChRs and a full agonist at α7 receptors (Mihalak et al. 2006). Varenicline is an FDA-approved smoking cessation treatment that reduces nicotine craving and withdrawal (Gonzales et al. 2006). A study with smokers suggests that varenicline may improve attention without affecting impulsive action (Lesage et al. 2020). Mecamylamine is a non-selective nAChR antagonist that has been shown to attenuate the behavioral and reinforcing effects of nicotine (Lundahl et al. 2000; Nickell et al. 2013). Several studies have indicated that mecamylamine may aid in smoking cessation, particularly when combined with nicotine replacement therapy (NRT)(Lancaster and Stead 1998; Rose et al. 1994). Nicotine replacement is another FDA-approved treatment for smoking cessation (Stead et al. 2012). Therefore, we also investigated the effects of nicotine pretreatment on responding for nicotine and impulsive action during the Go/No-Go test. By assessing the effects of these compounds on performance in the Go/No-Go task, we aimed to determine how modulation of nAChR signaling affects impulsive action after nicotine self-administration.

## 2. Material and methods

### 2.1. Animals

Adult male (220–280 g, 8-9 weeks of age; N=12) and female (180–225 g, 8-9 weeks of age; N=12) Wistar rats were purchased from Charles River (Raleigh, NC). The rats were housed with a rat of the same sex in a climate-controlled vivarium on a reversed 12 h light-dark cycle (light off at 7 AM). The rats were handled for 2-3 min per day for several days before surgery. Following jugular catheter implantation, rats were singly housed for the remainder of the study. Food and water were available ad libitum in the home cage throughout the study. The experimental protocols were approved by the University of Florida Institutional Animal Care and Use Committee (IACUC). All experiments were performed in accordance with relevant IACUC guidelines and regulations and in compliance with ARRIVE guidelines 2.0 (Animal Research: Reporting of *In Vivo* Experiments).

### 2.2. Drugs

For intravenous self-administration, (-)-nicotine hydrogen tartrate (NIDA Drug Supply Program) was dissolved in sterile saline (0.9 % sodium chloride), and the pH was adjusted to 7.2 ± 0.2 using 1M NaOH. Rats self-administered nicotine at a dose of 0.03 mg/kg/inf (expressed as base) in a volume of 0.1 ml/inf. For drug treatments, (-)-nicotine hydrogen tartrate, varenicline tartrate (Tocris bioscience, Minneapolis, MN), and mecamylamine hydrochloride (NIDA Drug Supply Program) were dissolved in sterile saline and administered subcutaneously (SC) in a volume of 1 ml/kg body weight. Nicotine doses are expressed as base, while varenicline and mecamylamine doses are expressed as salt.

### 2.3. Experimental design

A schematic overview of the experimental timeline is presented in Figure 1. Rats were surgically implanted with catheters in the jugular vein and allowed a minimum 7-day recovery period. Following recovery, male (N=12) and female (N=12) rats were trained to self-administer nicotine during fifteen 2 h sessions. This was followed by 41 days of nicotine self-administration under a Go/No-Go training paradigm. After completion of the Go/No-Go training sessions, the effects of varenicline, nicotine, and mecamylamine on nicotine self-administration in the Go/No-Go task were evaluated. In addition, a separate comparison examined responding in rats self-administering saline under the Go/No-Go schedule.

**Fig. 1.**
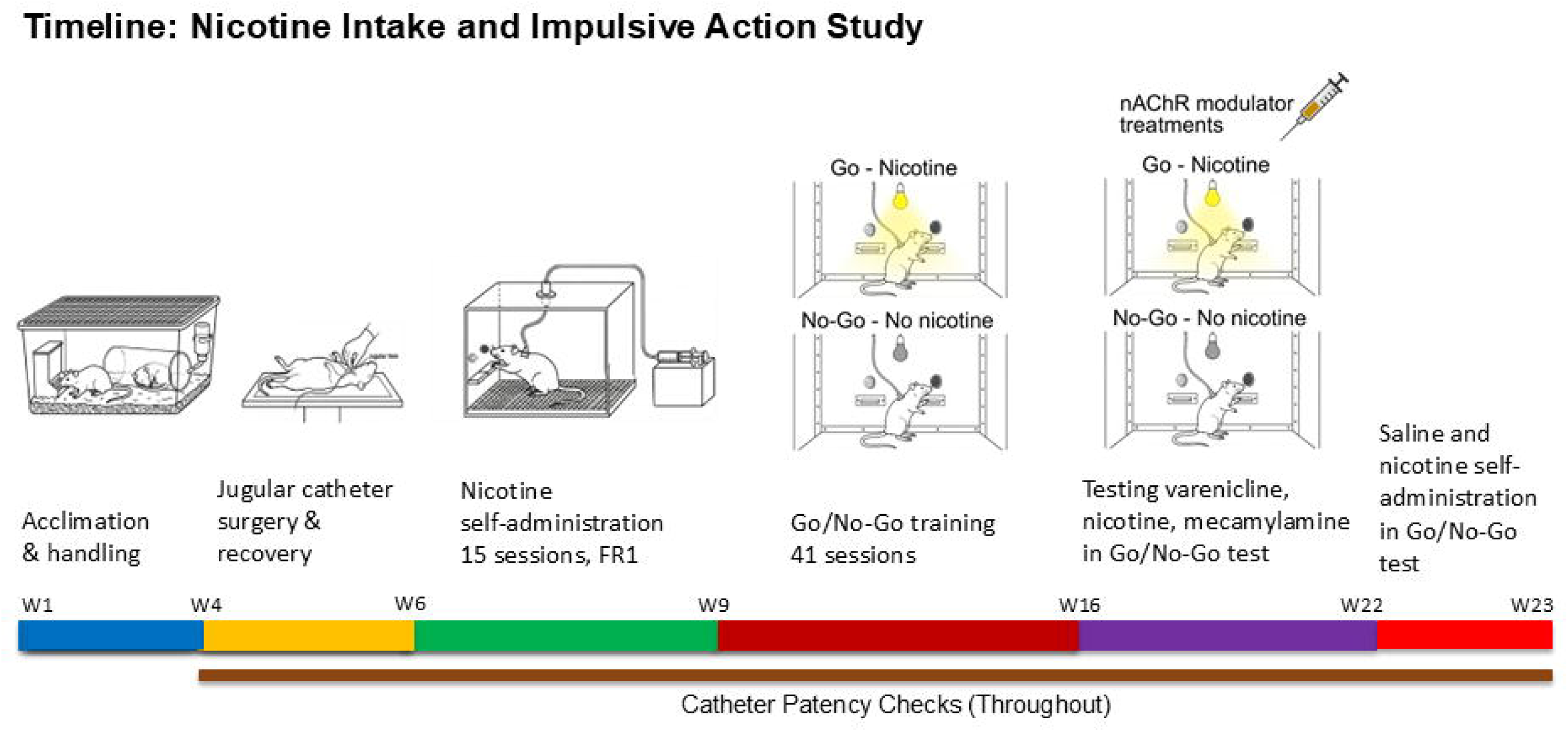
Schematic overview of the experimental timeline for the Go/No-Go nicotine self-administration study. Male and female rats were acclimated and handled prior to implantation of the jugular catheter followed by a minimum 7-day recovery period. Rats then acquired intravenous nicotine self-administration during fifteen 2-h sessions under an FR1 schedule. This was followed by 41 sessions of nicotine self-administration under a Go/No-Go training paradigm with alternating Go and No-Go periods. After completion of the Go/No-Go training, the effects of varenicline, nicotine, and mecamylamine on performance in the Go/No-Go task were assessed. In a separate comparison, rats self-administered saline under the Go/No-Go schedule to allow comparison with nicotine self-administration. Catheter patency was monitored throughout the study. Males, N=12; Females, N=12.

### 2.4. Intravenous catheter implantation

The catheters were implanted as described before (Chellian et al. 2024a; b). The rats were anesthetized with an isoflurane-oxygen vapor mixture (1-3%) and prepared with a catheter in the right jugular vein. The catheters consisted of polyurethane tubing (length 10 cm, inner diameter 0.64 mm, outer diameter 1.0 mm, model 3Fr, Instech Laboratories, Plymouth Meeting, PA). The right jugular vein was isolated, and the catheter was inserted 2.9 cm for males and 2.5 cm for females. The tubing was then tunneled subcutaneously and connected to a vascular access button (Instech Laboratories, Plymouth Meeting, PA). The button was exteriorized through a 1-cm incision between the scapulae. During the 7-day recovery period, the rats received daily infusions of the antibiotic Gentamycin (4 mg/kg, IV, Sigma-Aldrich, St. Louis, MO). A sterile heparin solution (0.1 ml, 50 U/ml) was flushed through the catheter before and after administering the antibiotic and after nicotine self-administration. After flushing the catheter, 0.05 ml of a sterile heparin/glycerol lock solution (500 U/ml, Instech Laboratories, Plymouth Meeting, PA) was infused into the catheter. The animals received carprofen (5 mg/kg, SC) daily for 72 hours after the surgery.

### 2.5. Acquisition of nicotine self-administration

Following recovery from surgery, rats were trained to acquire nicotine self-administration (0.03 mg/kg/infusion) under a fixed ratio 1 (FR1) schedule with a 10-second time-out (TO) in sound- and light-attenuated operant chambers (Med Associates, St. Albans, VT). Rats self-administered nicotine during daily 2 h sessions for 15 sessions. Nicotine self-administration sessions were conducted five days per week for three weeks. Responding on the active lever resulted in the delivery of a nicotine infusion (0.1 ml infused over a 6.5-s period). The infusion was paired with a cue light above the active lever, which remained illuminated throughout the time-out period. Responding on the inactive lever was recorded but did not have scheduled consequences. Both levers were retracted during the time-out (TO) period.

### 2.6. Go/No-Go nicotine self-administration task

The Go/No-Go self-administration task was conducted as previously described by Zapata et al. using cocaine (Zapata et al. 2017). Following the acquisition of nicotine self-administration, rats were trained to self-administer nicotine under the Go/No-Go schedule. Each session lasted 2 h and consisted of six alternating 20-minute intervals: three Go periods (nicotine available; 0–20 min, 40–60 min, and 80–100 min) and three No-Go periods (nicotine unavailable; 20–40 min, 60–80 min, and 100–120 min). Sessions always began with a Go period for all rats. Nicotine availability during the Go periods was signaled by illumination of the house light. During these periods, responding on the active lever resulted in an infusion of nicotine (0.03 mg/kg/infusion; 0.1 ml infused over 6.5 seconds) under a fixed ratio 5 (FR5) schedule with a 10-s timeout (TO). Each infusion was paired with a cue light above the active lever, which remained illuminated throughout the timeout period. The house light was turned off during the timeout period and turned on afterward to signal continued nicotine availability. Responses on the inactive lever were recorded but had no programmed consequences. Both levers were retracted during the timeout period. During the No-Go periods, the house light remained off to indicate that nicotine was unavailable. Lever presses were recorded during this period, but neither the active nor inactive lever produced any scheduled consequences. We assessed impulsive action using this task, specifically measuring the failure to inhibit responding during the No-Go period (commission errors). This task is a well-established paradigm for evaluating motor inhibition (Bari and Robbins 2013). Impulsive responding was quantified as the percentage of active lever responses emitted during No-Go periods relative to total active lever responses across both Go and No-Go periods, calculated as: [No-Go active lever responses / (Go active lever responses + No-Go active lever responses) × 100 (Zapata and Lupica 2021). This measure indexes commission errors, reflecting impaired inhibitory control when nicotine was unavailable. Because nicotine infusions during Go periods were obtained under an FR5 schedule, overall active lever responding was higher during Go intervals. Consequently, relying on raw counts of No-Go responses may misrepresent impulsivity, as such differences can arise from variations in overall response output or reinforcement contingencies rather than true deficits in inhibitory control. Expressing commission errors as a proportion of total active-lever responses accounts for these differences and provides a rate-independent measure of inhibitory control. Go/No-Go training was conducted for 41 consecutive sessions. The first 20 sessions were conducted five days per week, while the remaining 21 sessions were conducted seven days per week.

### 2.7. Varenicline, nicotine, and mecamylamine treatment on Go/No-Go nicotine self-administration task

Following the Go/No-Go training sessions, the effects of varenicline, nicotine, and mecamylamine were assessed on nicotine self-administration behavior within the Go/No-Go task. Each drug was administered 15 minutes prior to the Go/No-Go nicotine self-administration (0.03 mg/kg/infusion) session. Varenicline (0, 0.1, 0.3, 1, and 3 mg/kg, SC), nicotine (0, 0.1, 0.3, and 0.6 mg/kg, SC), and mecamylamine (0, 0.5, 1, and 2 mg/kg, SC) were administered using a Latin square design. There was at least a 48-hour interval between successive test sessions for each drug (varenicline, nicotine, or mecamylamine). Daily Go/No-Go nicotine self-administration sessions were conducted between the treatment sessions. The treatments with nicotine began 72 hours after the last varenicline session, and the treatments with mecamylamine began 72 hours after the last nicotine session. Go/No-Go nicotine self-administration sessions were conducted seven days per week.

### 2.8. Saline versus nicotine self-administration in the Go/No-Go task

After completion of all drug treatment experiments, a separate comparison was conducted to evaluate responding for saline versus nicotine under the Go/No-Go schedule. Rats continued to undergo daily Go/No-Go nicotine self-administration and ninety-six hours after the final drug treatment session, rats were assigned to a within-subject crossover design. During the first test phase, half of the animals self-administered saline, while the remaining animals self-administered nicotine (0.03 mg/kg/infusion) under identical Go/No-Go task conditions. Following this initial test, animals underwent two standard Go/No-Go nicotine self-administration sessions to re-establish baseline responding before crossing over to the alternate condition. Animals that initially self-administered saline subsequently self-administered nicotine, and animals that initially self-administered nicotine subsequently self-administered saline. Go/No-Go nicotine self-administration sessions were conducted seven days per week.

### 2.8. Catheter patency test

Catheter patency was assessed during the self-administration period by administering 0.2 ml of the ultra-short-acting barbiturate Brevital (1% methohexital sodium). A rapid loss of muscle tone was considered indicative of a patent catheter. Rats that did not exhibit this response were excluded from subsequent analyses. One male and one female rat failed the Brevital test during the Go/No-Go training period and were excluded. Additionally, two male rats did not respond to Brevital during the nicotine treatment phase and were excluded.

### 2.9. Statistics

Data were analyzed with SPSS Statistics version 31 and GraphPad Prism version 10.5. The figures were generated using GraphPad Prism version 10.5. Lever pressing, percentage of active lever responses, nicotine intake, and infusion latency data were analyzed using two-way or three-way analysis of variance (ANOVA) with repeated measures where appropriate. The factors included treatment, sex, session, and lever (active vs. inactive). Significant interactions and main effects were further examined using the Bonferroni post hoc test. The alpha level for statistical significance was set at p < 0.05.

## 3. Results

### 3.1. Acquisition of nicotine self-administration

Male and female rats acquired nicotine self-administration over 15 sessions. During this period, both male and female rats responded more on the active lever than the inactive lever (Fig. S1A, Lever F1,22=31.191, P < 0.001; Sex F1,22=0.033, NS; Lever × Sex F1,22=1.209, NS). Inactive lever responding was stable, while active lever responding increased over time and then stabilized in both males and females (Fig. S1A, Session F14,308=4.753, P < 0.001; Session × Sex F14,308=0.787, NS; Lever × Session F14,308=2.717, P < 0.001; Lever × Session × Sex F14,308=0.771, NS). Nicotine intake initially increased and then stabilized in males and females (Fig. S1B, Session F14,308=12.583, P < 0.001; Session × Sex F14,308=1.698, NS; Sex F1,22=0.507, NS). Furthermore, the latency to the first nicotine infusion exhibited a sex-dependent pattern: in males, the latency initially increased before decreasing and stabilizing, whereas in females, it decreased early and remained stable thereafter (Fig. S1C, Session F14,308=7.266, P < 0.001; Sex F1,22=2.571, NS; Session × Sex F14,308=1.861, P < 0.05). The post hoc analysis revealed that females had significantly shorter first infusion latency than males on session 2 (Fig. S1C). In addition, the post hoc analysis showed that the latency to the first infusion in males was significantly longer on session two and significantly shorter from session six onward, compared to session one (Fig. S1C).

### 3.2. Go/No-Go training

After the acquisition phase, male and female rats were trained to self-administer nicotine in a Go/No-Go task for 41 sessions.

#### Go period

During the Go period, active lever responses were higher than inactive lever responses, with no effect of sex (Fig. 2A, Lever F1,20=137.087, P < 0.001; Sex F1,20=0.276, NS; Lever × Sex F1,20=0.078, NS). Active lever responding increased during the initial sessions and then remained relatively stable in both males and females, while inactive lever responses were relatively stable (Fig. 2A, Session F40,800=4.515, P < 0.001; Session × Sex F40,800=1.042, NS; Lever × Session F40,800=2.384, P < 0.001; Lever × Session × Sex F40,800=1.35, NS). Nicotine intake quickly increased and then stabilized in both males and females (Fig. 2B, Session F40,80=5.008, P < 0.001; Session × Sex F40,80=1.197, NS; Sex F1,20=0.272, NS).

**Fig. 2.**
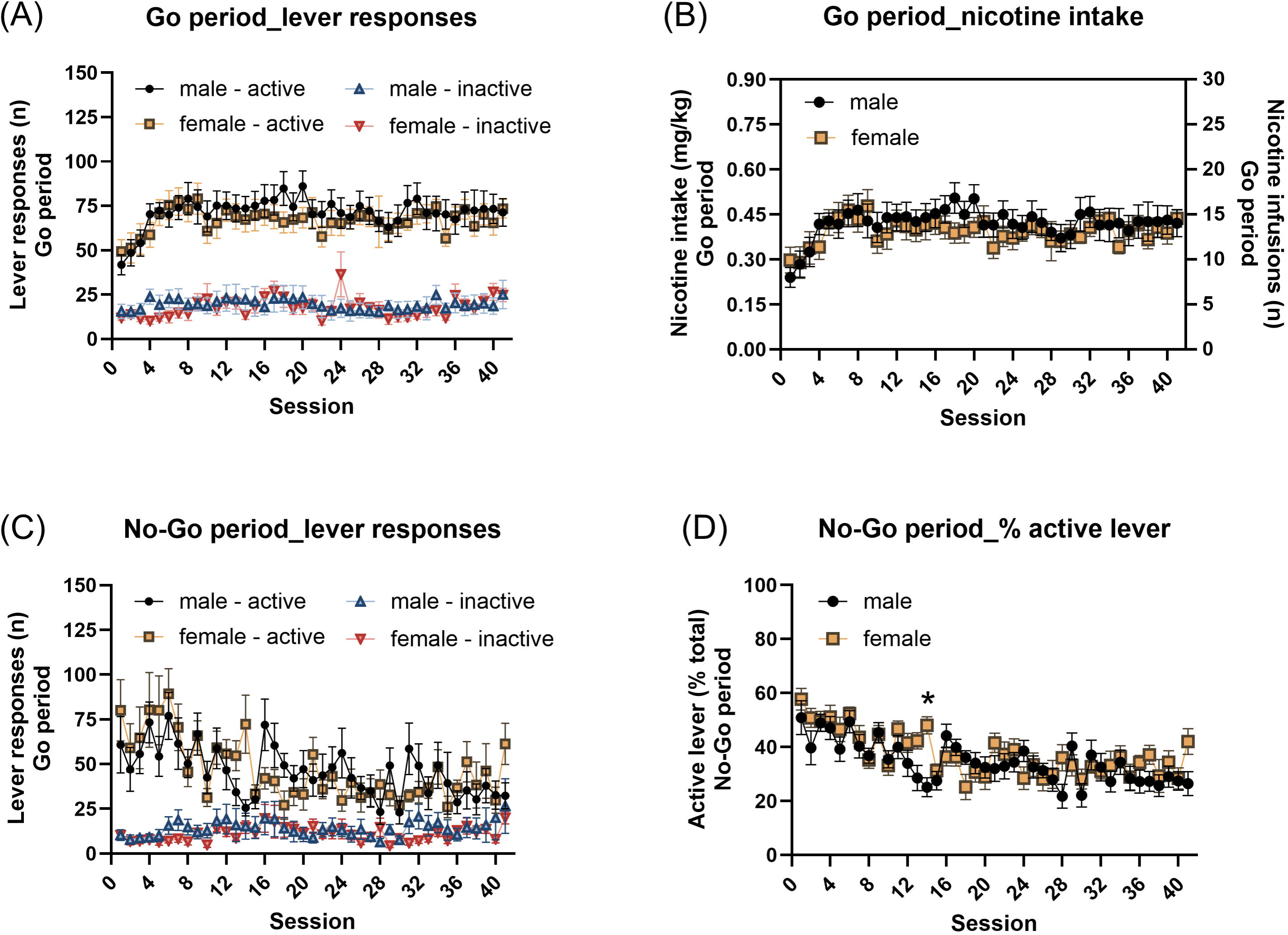
Nicotine self-administration and responding during the Go/No-Go training phase. (A) Active and inactive lever responses during the Go periods across the 41 training sessions. Active lever responding increased and stabilized, remaining significantly higher than inactive lever responding. (B) Nicotine intake (mg/kg) during the Go periods, which increased and stabilized over the training sessions. (C) Active and inactive lever responses during the No-Go periods. Active lever responding decreased over time, consistent with the acquisition of inhibitory control, but remained significantly higher than inactive lever responding. (D) Percentage of active lever responses during the No-Go periods across training sessions, which decreased over time in both sexes. Males, N=11; Females, N=11. * p < 0.05. Data are expressed as mean + SEM.

#### No-Go period

During the No-Go period, the rats responded more on the active lever than the inactive lever, with no effect of sex (Fig. 2C, Lever F1,20=102.625, P < 0.001; Sex F1,20=0.024, NS; Lever × Sex F1,20=0.43, NS). Lever-pressing patterns changed significantly over time and differed between sexes (Session F40,800=4.313, P < 0.001; Session × Sex F40,800=1.745, P < 0.01; Lever × Session F40,800=5.242, P < 0.001; Lever × Session × Sex F40,800=1.768, P < 0.01). The post hoc analysis did not reveal significant sex differences in either active or inactive lever responses when comparing the same session between males and females. However, the post hoc analysis showed that the active lever responses significantly decreased from session 14 onward in males and from session 8 onward in females compared to the session 1 within the same sex (Table S1). In contrast, no significant changes in inactive lever responses were observed across sessions in either sex when compared to their respective session 1.

The percentage of active lever responses during the No-Go period decreased over time and this pattern was affected by sex (Fig. 2D, Session F40,80=8.144, P < 0.001; Sex F1,20=0.864, NS; Session × Sex F40,80=2.076, P < 0.01). The post hoc analysis showed that the females had a significantly higher percentage of active lever responses than males during session 14 (Fig. 2D). In addition, the post hoc analysis revealed that the percentage active lever responses significantly decreased from session 12 onward in males and from session 8 onward in females compared to the session 1 within the same sex (Table S1).

### 3.3. Go/No-Go task: varenicline treatment

#### Go period

Varenicline treatment decreased both active and inactive lever responses during the Go period, and these effects were not affected by sex (Fig. 3A, Active lever: Varenicline treatment F4,80=48.664, P < 0.001; Varenicline treatment × Sex F4,80=0.753, NS; Sex F1,20=0.223, NS; Fig. 3C, Inactive lever: Varenicline treatment F4,80=9.326, P < 0.001; Varenicline treatment × Sex F4,80=0.147, NS; Sex F1,20=0.069, NS). Furthermore, varenicline reduced nicotine intake in both males and females (Fig. 3B, Varenicline treatment F4,80=46.811, P < 0.001; Varenicline treatment × Sex F4,80=0.397, NS; Sex F1,20=0.275, NS).

**Fig. 3.**
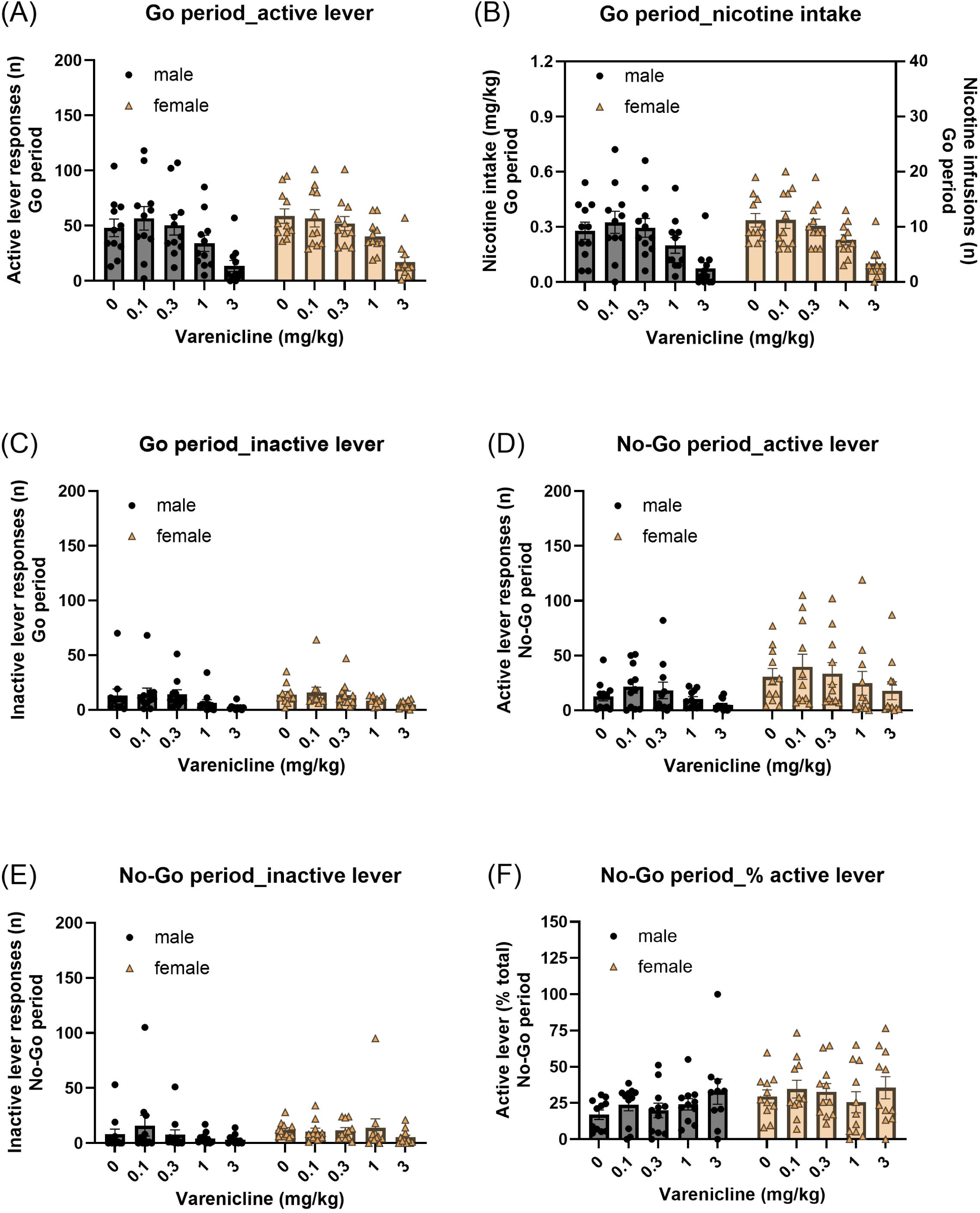
Effects of varenicline treatment on performance in the nicotine Go/No-Go task. Effects of varenicline (0, 0.1, 0.3, 1, and 3 mg/kg) on (A) active lever responses, (B) nicotine intake, and (C) inactive lever responses during the Go periods. Varenicline dose-dependently reduced responding and intake during Go periods. (D) Active lever responses, (E) inactive lever responses, and (F) the percentage of active lever responses during the No-Go periods. Varenicline decreased active lever responses during No-Go periods but did not alter the percentage of active responses (response allocation), indicating a nonspecific reduction in responding rather than specific improvement in inhibitory control. Males, N=11; Females, N=11. Data are expressed as mean + SEM.

#### No-Go period

Varenicline treatment decreased active lever responses during the No-Go period in both male and female rats (Fig. 3D, Varenicline treatment F4,80=5.086, P < 0.01; Varenicline treatment × Sex F4,80=0.112, NS; Sex F1,20=3.007, NS). Inactive lever responses were not significantly affected by varenicline treatment (Fig. 3E, Varenicline treatment F4,80=1.631, NS; Treatment × Sex F4,80=1.089, NS; Sex F1,20=0.327, NS). Similarly, varenicline treatment did not affect the percentage of active lever responses during the No-Go period (Fig. 3F, Varenicline treatment F4,80=1.806, NS; Varenicline treatment × Sex F4,80=0.762, NS; Sex F1,20=1.975, NS).

### 3.4. Go/No-Go task: nicotine treatment

#### Go period

Pretreatment with nicotine decreased active and inactive lever responses during the Go period, with no effects of sex (Fig. 4A, Active lever: Nicotine treatment F3,54=41.204, P < 0.001; Nicotine treatment × Sex F3,54=0.501, NS; Sex F1,18=0.404, NS; Fig. 4C, Inactive lever: Nicotine treatment F3,54=5.322, P < 0.01; Nicotine treatment × Sex F3,54=0.035, NS; Sex F1,18=0.04, NS). In addition, nicotine intake was decreased after nicotine treatment in both males and females (Fig. 4B, Nicotine treatment F3,54=41.944, P < 0.001; Nicotine treatment × Sex F3,54=0.69, NS; Sex F1,18=0.371, NS).

**Fig. 4.**
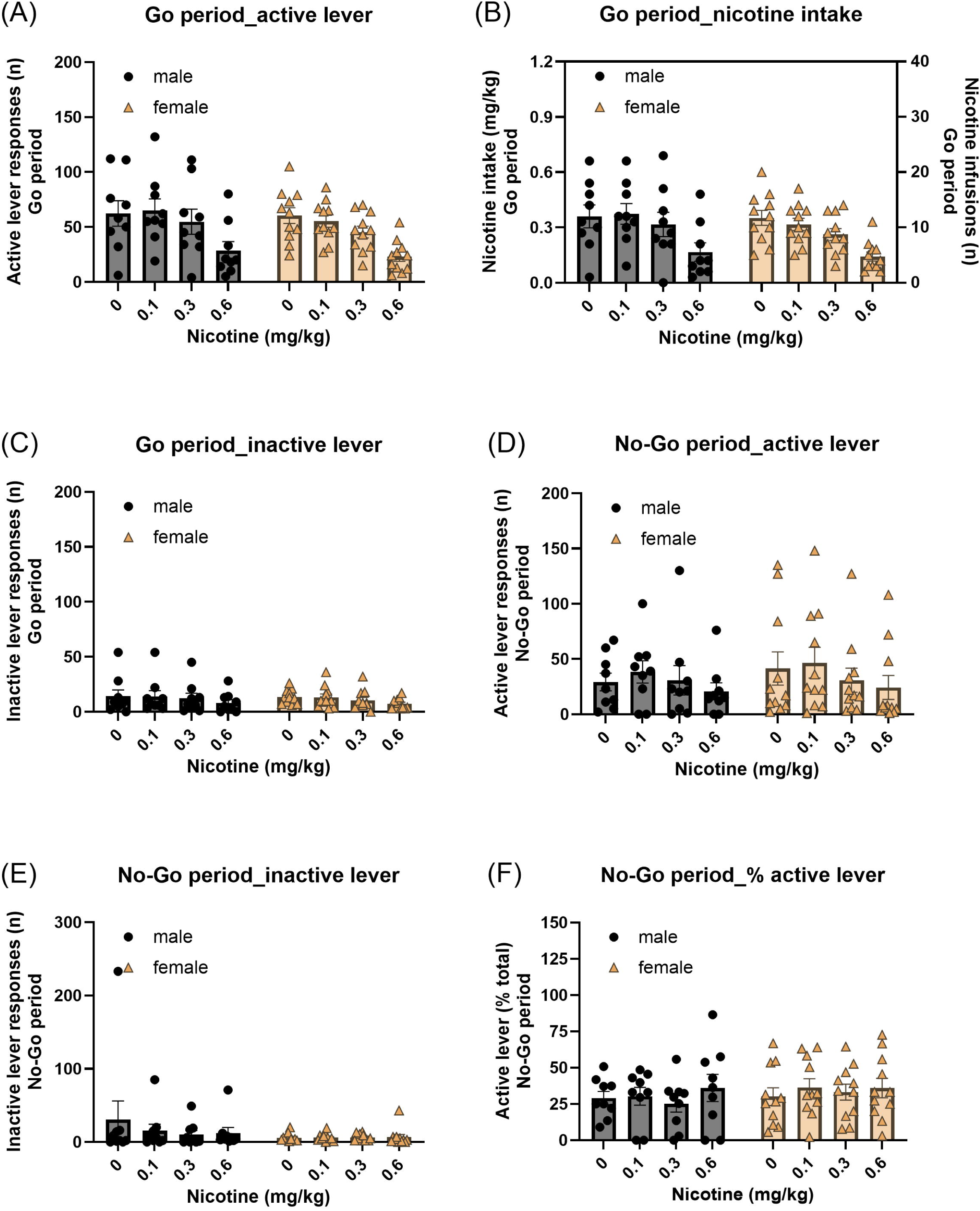
Effects of nicotine pretreatment on performance in the nicotine Go/No-Go task. Effects of nicotine pretreatment (0, 0.1, 0.3, and 0.6 mg/kg) on (A) active lever responses, (B) nicotine intake, and (C) inactive lever responses during the Go periods. Nicotine pretreatment decreased active responding and intake during Go periods. (D) Active lever responses, (E) inactive lever responses, and (F) the percentage of active lever responses during the No-Go periods. Similar to varenicline, nicotine decreased active lever responses during No-Go periods but did not alter the percentage of active responses, suggesting nonspecific suppression of behavior rather than enhanced response inhibition. Males, N=9; Females, N=11. Data are expressed as mean + SEM.

#### No-Go period

During the No-Go period, nicotine treatment decreased active lever responding in both males and females (Fig. 4D, Nicotine treatment F3,54=3.471, P < 0.05; Nicotine treatment × Sex F3,54=0.351, NS; Sex F1,18=0.161, NS), while inactive lever responses were not affected (Fig. 4E, Nicotine treatment F3,54=1.081, NS; Nicotine treatment × Sex F3,54=1.223, NS; Sex F1,18=1.056, NS). Furthermore, the percentage of active lever responses was not affected by nicotine treatment in either males or females (Fig. 4F, Nicotine treatment F3,54=0.971, NS; Nicotine treatment × Sex F3,54=0.352, NS; Sex F1,18=0.286, NS).

### 3.5. Go/No-Go task: mecamylamine treatment

#### Go period

Mecamylamine treatment did not affect active lever responses during the Go period, and no sex differences were observed (Fig. 5A, Mecamylamine treatment F3,54=2.08, NS; Mecamylamine treatment × Sex F3,54=0.793, NS; Sex F1,18=1.512, NS). However, inactive lever responses were reduced by mecamylamine treatment, independent of sex (Fig. 5C, Mecamylamine treatment F3,54=5.233, P < 0.01; Mecamylamine treatment × Sex F3,54=1.182, NS; Sex F1,18=1.429, NS). Mecamylamine produced a trend toward reduced nicotine intake, however, this effect did not reach statistical significance (Fig. 5B, Mecamylamine treatment F3,54= 2.749, P = 0.052; Mecamylamine treatment × Sex F3,54=0.644, NS; Sex F1,18=1.86, NS).

**Fig. 5.**
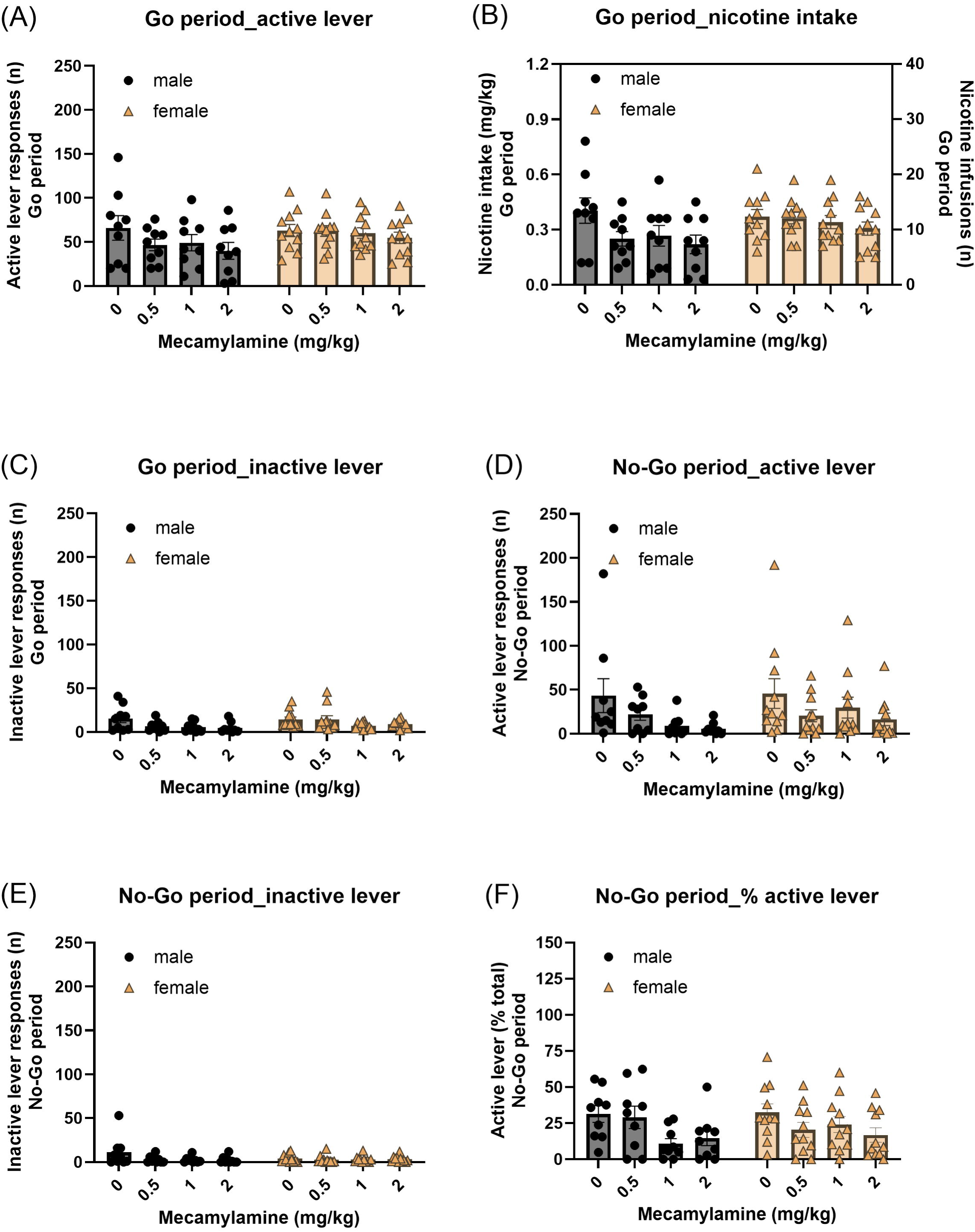
Effects of mecamylamine treatment on performance in the nicotine Go/No-Go task. Effects of mecamylamine (0, 0.5, 1, and 2 mg/kg) on (A) active lever responses, (B) nicotine intake, and (C) inactive lever responses during the Go periods. Mecamylamine reduced did not significantly alter active lever responses and nicotine intake during Go periods. (D) Active lever responses, (E) inactive lever responses, and (F) the percentage of active lever responses during the No-Go periods. Mecamylamine significantly reduced both active lever responding and the percentage of active responses during No-Go periods, consistent with a selective reduction in drug-directed response bias. Males, N=9; Females, N=11. Data are expressed as mean + SEM.

#### No-Go period

During the No-Go period, mecamylamine treatment decreased both active and inactive lever responses in both males and females (Fig. 5D, Active lever: Mecamylamine treatment F3,54=5.028, P < 0.01; Mecamylamine treatment × Sex F3,54=0.63, NS; Sex F1,18=0.483, NS; Fig. 5E, Inactive lever: Mecamylamine treatment F3,54=3.532, P < 0.05; Mecamylamine treatment × Sex F3,54=2.103, NS; Sex F1,18=0.477, NS). Mecamylamine administration also decreased the percentage of active lever responses during the No-Go period (Fig. 5F, Mecamylamine treatment F3,54=4.114, P < 0.05; Mecamylamine Treatment × Sex F3,54=2.35, NS; Sex F1,18=0.177, NS).

### 3.6. Go/No-Go task: Saline and nicotine self-administration

#### Go period

During the Go period, active lever responses and infusions were similar between the Nicotine Group and Saline Group, with no effects of sex (Fig. 6A, Active lever: Group F1,18=0.166, NS; Group × Sex F1,18=0.547, NS; Sex F1,18=0.15, NS; Fig. 6B, Infusions: Group F1,18=0.058, NS; Group × Sex F1,18=0.675, NS; Sex F1,18=0.215, NS). However, inactive lever responses during the Go period were significantly higher in the Nicotine Group compared to the Saline Group, regardless of sex (Fig. 6C, Group F1,18=18.917, P < 0.001; Group × Sex F1,18=0.029, NS; Sex F1,18=1.12, NS).

**Fig. 6.**
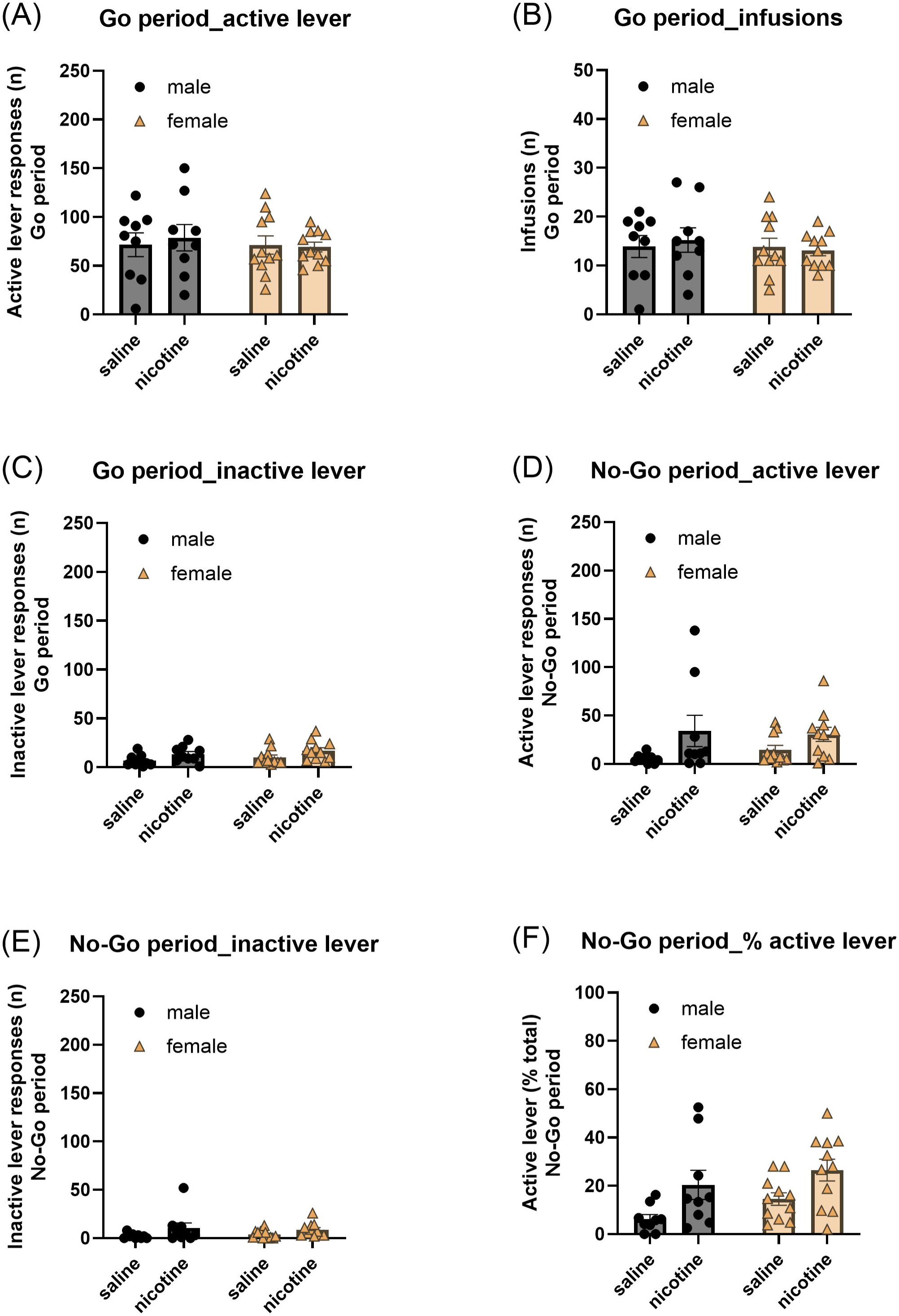
Comparison of impulsive action in rats self-administering nicotine versus saline. (A) Active lever responses, (B) total infusions, and (C) inactive lever responses during the Go periods for rats self-administering nicotine vs. saline. (D) Active lever responses, (E) inactive lever responses, and (F) percentage of active lever responses during the No-Go periods. Rats self-administering nicotine displayed significantly higher active lever responding and a higher percentage of active responses during No-Go periods compared to saline controls, consistent with increased impulsive action. Males, N=9; Females, N=11. Data are expressed as mean + SEM.

#### No-Go period

During the No-Go period, rats in the Nicotine Group had more active and inactive lever responses than those in the Saline Group, with no significant sex differences (Fig 6D, Active lever: Group F1,18=8.022, P < 0.05; Group × Sex F1,18=0.677, NS; Sex F1,18=0.087, NS; Fig 6E, Inactive lever: Group F1,18=5.403, P < 0.05; Group × Sex F1,18=0.35, NS; Sex F1,18=0.003, NS). Similarly, the percentage of active lever responses during the No-Go period was higher in the Nicotine Group compared to the Saline Group, independent of sex (Fig 6F, Group F1,18=14.012, P < 0.001; Group × Sex F1,18=0.091, NS; Sex F1,18=2.522, NS).

## Discussion

In this study we investigated whether nAChR modulators affect impulsive action in a nicotine self-administration Go/No-Go task in male and female rats. The present study demonstrates that chronic nicotine self-administration increases impulsive action in rats. During the Go/No-Go training period, responses on the active lever increased during Go periods and decreased during No-Go periods, indicating effective acquisition of both the ‘go’ response and inhibitory control. The rats also displayed higher active than inactive lever responses during No-Go periods, indicating high levels of impulsive action under conditions requiring inhibitory control Acute pretreatment with varenicline or nicotine reduced active lever responding during both Go and No-Go periods, but did not alter response allocation, indicating that these reductions reflect nonspecific decreases in responding rather than improvements in inhibitory control. The non-selective nAChR antagonist mecamylamine did not affect active lever responses during the Go period but decreased active lever responding during No-Go periods. Finally, substituting nicotine with saline infusions did not alter active lever responding during Go periods, but led to a decrease in active lever responding during No-Go periods compared to rats self-administering nicotine, confirming that chronic nicotine self-administration enhances impulsivity. These findings emphasize the utility of the Go/No-Go paradigm as a translational model to investigate nicotine-induced impairments in inhibitory control and to identify potential therapeutic interventions targeting impulsive behavior associated with tobacco use disorder.

In the present study, we found that nicotine self-administration increased responding during the No-Go period and increased the percentage of active responses during the No-Go period compared with the saline self-administration, indicating increased impulsive action. This observation is in line with a previous Go/No-Go study that investigated the effects of noncontingently administered nicotine on impulsive behavior (Kolokotroni et al. 2011). Kolokotroni et al., investigated the effects of nicotine in a symmetrically reinforced Go/No-Go task and a systematic delayed reward task (Kolokotroni et al. 2011). Acute nicotine administration induced a dose-dependent increase in both impulsive choice and increased responding during No-Go trials indicating increased impulsive action. These studies with the Go/No-Go task indicate that both acute and chronic treatment with nicotine increases impulsivity. Studies using the 5-Choice Serial Reaction Time Task (5-CSRTT), which is a well-established assay for assessing visuospatial attention and impulsive action, also indicate that acute and chronic treatment with nicotine affects impulsive behavior (Asinof and Paine 2014). For example, Day et al. reported that repeated nicotine exposure decreased response latencies and increased inappropriate responding, indicative of heightened impulsivity in rats previously exposed to nicotine (Day et al. 2007). Similarly, Blondel et al. found that acute and chronic nicotine treatment increased anticipatory responding in rats (Blondel et al. 2000).

In the present study we also investigated the effects of the nAChR modulators nicotine, mecamylamine, and varenicline on responding in the Go/No-Go paradigm. Pretreatment with nicotine decreased nicotine intake during the Go period. Consistent with previous findings, pretreatment with nicotine dose-dependently reduced nicotine self-administration, likely due to nAChR occupancy or desensitization that diminishes the reinforcing effects of nicotine (Corrigall and Coen 1989; Dani 2015; Green et al. 2000). Pretreatment with nicotine also led to a decrease in active lever responses during the No-Go period; however, because the percentage of active responses was unchanged, this decrease likely reflects reduced overall responding rather than enhanced inhibitory control. Pretreatment with nicotine reduced nicotine self-administration, but this decrease in intake did not translate into reduced impulsive action, as response allocation during No-Go periods remained unchanged. Previous studies indicate that nicotine-induced impulsive action is primarily mediated by activation of nAChRs (Blondel et al. 2000; Kolokotroni et al. 2011). In the present paradigm, both nicotine pretreatment and self-administered nicotine likely engaged the same nAChR-dependent mechanisms during task performance, resulting in sustained receptor activation despite reduced nicotine intake. As a result, nicotine pretreatment did not change nicotine-associated impulsive action, consistent with the maintenance of impaired inhibitory control driven by ongoing nAChR activation. These findings indicate that reductions in nicotine reinforcement can occur independently of improvements in inhibitory control and highlight the importance of nAChR signaling in nicotine-associated impulsivity in the Go/No-Go self-administration task.

In the present study, we also investigated the effects of the nAChR antagonist mecamylamine on nicotine intake during the Go period and impulsive responding during the No-Go period. Mecamylamine did not significantly affect active lever presses or nicotine intake. Conflicting findings have been reported regarding the effects of mecamylamine on responding for nicotine (Chellian et al. 2024a; b; Corrigall and Coen 1989). Mecamylamine has been shown to decrease nicotine in rats with a relatively short history of nicotine intake (Corrigall and Coen 1989). In contrast, mecamylamine increases nicotine intake in nicotine dependent rats with a long history of nicotine intake (Chellian et al. 2024b). In our study, treatment with mecamylamine also decreased responding during the No-Go period; combined with the observed reduction in the percentage of active responses, this indicates that mecamylamine reduced impulsive action in rats that self-administered nicotine. This finding is consistent with previous reports that nicotine-induced impulsivity is attenuated by mecamylamine pretreatment (Blondel et al. 2000; Kolokotroni et al. 2011). These effects likely reflect nAChR blockade within pathways mediating nicotine-induced impulsivity, confirming that the enhanced impulsive action observed in our paradigm is nAChR-dependent. Although we did not test mecamylamine in saline self-administering rats in this study, prior work suggests that mecamylamine does not alter impulsive behavior in drug-naive animals tested in symmetrically reinforced Go/No-Go tasks, systematic delayed reward tasks, or the 5-CSRTT (Blondel et al. 2000; Kolokotroni et al. 2011). Together, these findings support the conclusion that nicotine self-administration-induced impulsivity is specifically mediated via nAChR activation and demonstrate the utility of this model for investigating pharmacological modulation of drug-induced impulsive behavior.We also investigated the effects of varenicline on responding during the Go and No-Go period. Varenicline (a partial agonist at α4β2 nAChRs and a full agonist at α7 receptors) is an FDA approved smoking cessation drug and decreases nicotine withdrawal and intake in rats (Faessel et al. 2010; Igari et al. 2013). In our study, varenicline decreased active lever responses during the Go period and during the No-Go period. The observation that varenicline decreases responding during the Go period is in line with previous studies showing that varenicline decreases nicotine self-administration (George et al. 2011; O’Connor et al. 2010). This study also showed that varenicline decreases responding during the No-Go period; however, the lack of change in the percentage of active responses indicates that varenicline did not modify impulsive action but instead reduced responding nonspecifically. These results are consistent with prior work using the 3-choice serial reaction time tasks (3-CSRTT), which showed that varenicline pretreatment does not alter nicotine-induced impulsive behavior (Ohmura et al. 2017). Although varenicline was not tested in saline self-administration, previous studies suggest that it can induce impulsive responding in drug-naïve animals in both 3- and 5-CSRTT paradigms, an effect attributed to its partial agonism at α4β2 nAChRs (Ohmura et al. 2017; Wouda et al. 2011). Partial agonists bind to the same receptor sites as full agonists such as nicotine but produce a submaximal receptor response; this partial activation of nAChRs has been shown to increase impulsive action in models of inhibitory control. In our paradigm, varenicline’s acute partial activation of nAChRs likely reduced nicotine-maintained responding without affecting inhibitory control, consistent with the unchanged percentage of active responses during No-Go periods. Because varenicline competes with nicotine for nAChR binding yet only partially activates these receptors, the receptor signaling necessary for impulsive action remains engaged even as nicotine intake decreases. Together, these findings indicate that reductions in nicotine intake reflect decreased reinforcement rather than modulation of nicotine-induced impulsivity and further support the validity of the Go/No-Go self-administration model for investigating nAChR-mediated impulsive action.

The percentage of active responses during No-Go trials served as an index of response bias toward the nicotine-associated lever. Neither nicotine nor varenicline altered this percentage, indicating that α4β2* nAChR partial agonism did not modify response allocation. In other words, although these compounds decreased the overall responding, they did not change which lever the animals chose, indicating that bias toward the nicotine-paired lever remained intact. Importantly, if these reductions were driven by motor incapacity, we would expect a disruption in lever discrimination. Thus, the unchanged percentage of active responses indicates that nicotine and varenicline did not impair motor capacity, but rather reduced responding nonspecifically. By contrast, mecamylamine significantly reduced the percentage of active responses, reflecting a shift away from persistent responding on the nicotine-associated lever during periods requiring response inhibition. This selective reduction in drug-directed response bias suggests that nAChR antagonism reduces the tendency to select the nicotine-associated lever, rather than merely suppressing general responding. Collectively, these findings demonstrate that the percentage active response measure is sensitive to nicotine-related bias in action selection and that only mecamylamine altered this bias under No-Go conditions.

In conclusion, this study demonstrates that chronic nicotine self-administration produces an increase in impulsive responding in a Go/No-Go task, which is indicative of a deficit in inhibitory control. Our findings show that drugs targeting nAChRs affect performance in the Go/No-Go task. However, only mecamylamine reduced impulsive action as measured by the percentage of active responses; nicotine and varenicline reduced overall responding without altering response allocation. Furthermore, rats that self-administered saline showed lower impulsive responding during the No-Go period than rats self-administering nicotine, indicating that active nicotine intake drives the expression of impulsive behavior, rather than chronic exposure to nicotine producing a permanent deficit in inhibitory control. This work supports the utility of the Go/No-Go task as a valuable translational model for investigating the neurobiology of nicotine addiction and for developing novel therapeutics aimed at improving inhibitory control in people who use tobacco products.

## CRediT authorship contribution statement

**Adriaan W. Bruijnzeel:** Conceptualization, Supervision, Formal analysis, Writing – Original Draft, Visualization, Project administration, Funding acquisition.

**Guido Huisman:** Investigation, Project administration.

**Ranjithkumar Chellian:** Formal analysis, Investigation, Writing – review & editing, Visualization.

## Data Integrity and Sponsor Role

The authors confirm that they have had full access to all the data in the study and take responsibility for the integrity of the data and the accuracy of the data analysis. No sponsor was involved in the study design, data collection, or writing of the manuscript.

## Design and Analysis Transparency

The datasets generated and analyzed during the current study are available from the corresponding author on request.

