## Supplemental material for "Nicotine self-administration increases impulsive action: differential effects of nAChR modulators in a Go/No-Go task"

**Table S1: NoGo period lever responses(compared to session 1).**

| **Session** | **Fig. 2C: NoGo period: lever responses** | | **Fig. 2D: NoGo period: percentage active lever responses** | |
| --- | --- | --- | --- | --- |
|  | **males** | **females** | **males** | **females** |
| **2** | ns | ns | ns | ns |
| **3** | ns | ns | ns | ns |
| **4** | ns | ns | ns | ns |
| **5** | ns | ns | ns | ns |
| **6** | ns | ns | ns | ns |
| **7** | ns | ns | ns | ns |
| **8** | ns | ** | ns | *** |
| **9** | ns | ns | ns | ns |
| **10** | ns | *** | ns | *** |
| **11** | ns | ns | ns | ns |
| **12** | ns | ns | * | * |
| **13** | ns | ns | *** | ns |
| **14** | ** | ns | *** | ns |
| **15** | * | *** | *** | *** |
| **16** | ns | *** | ns | *** |
| **17** | ns | *** | ns | *** |
| **18** | ns | *** | ns | *** |
| **19** | ns | *** | * | *** |
| **20** | ns | *** | ** | *** |
| **21** | ns | ns | ** | * |
| **22** | ns | *** | ** | ** |
| **23** | ns | ** | * | ** |
| **24** | ns | *** | ns | *** |
| **25** | ns | *** | ** | *** |
| **26** | ns | *** | ** | *** |
| **27** | ns | *** | *** | *** |
| **28** | ** | *** | *** | *** |
| **29** | ns | *** | ns | *** |
| **30** | ** | *** | *** | *** |
| **31** | ns | *** | ns | *** |
| **32** | ns | *** | ** | *** |
| **33** | ns | *** | *** | *** |
| **34** | ns | * | * | *** |
| **35** | ns | *** | *** | *** |
| **36** | * | *** | *** | *** |
| **37** | ns | ns | *** | ** |
| **38** | * | *** | *** | *** |
| **39** | ns | ** | *** | *** |
| **40** | ns | *** | *** | *** |
| **41** | ns | ns | *** | ns |
| Bonferroni's multiple comparisons post hoc test: Asterisks indicate a significant difference from session 1 in rats of the same sex. * P < 0.05, ** P < 0.01, *** P< 0.001. Males, N=11; Females, N=11. | | | | |

**Figure S1**

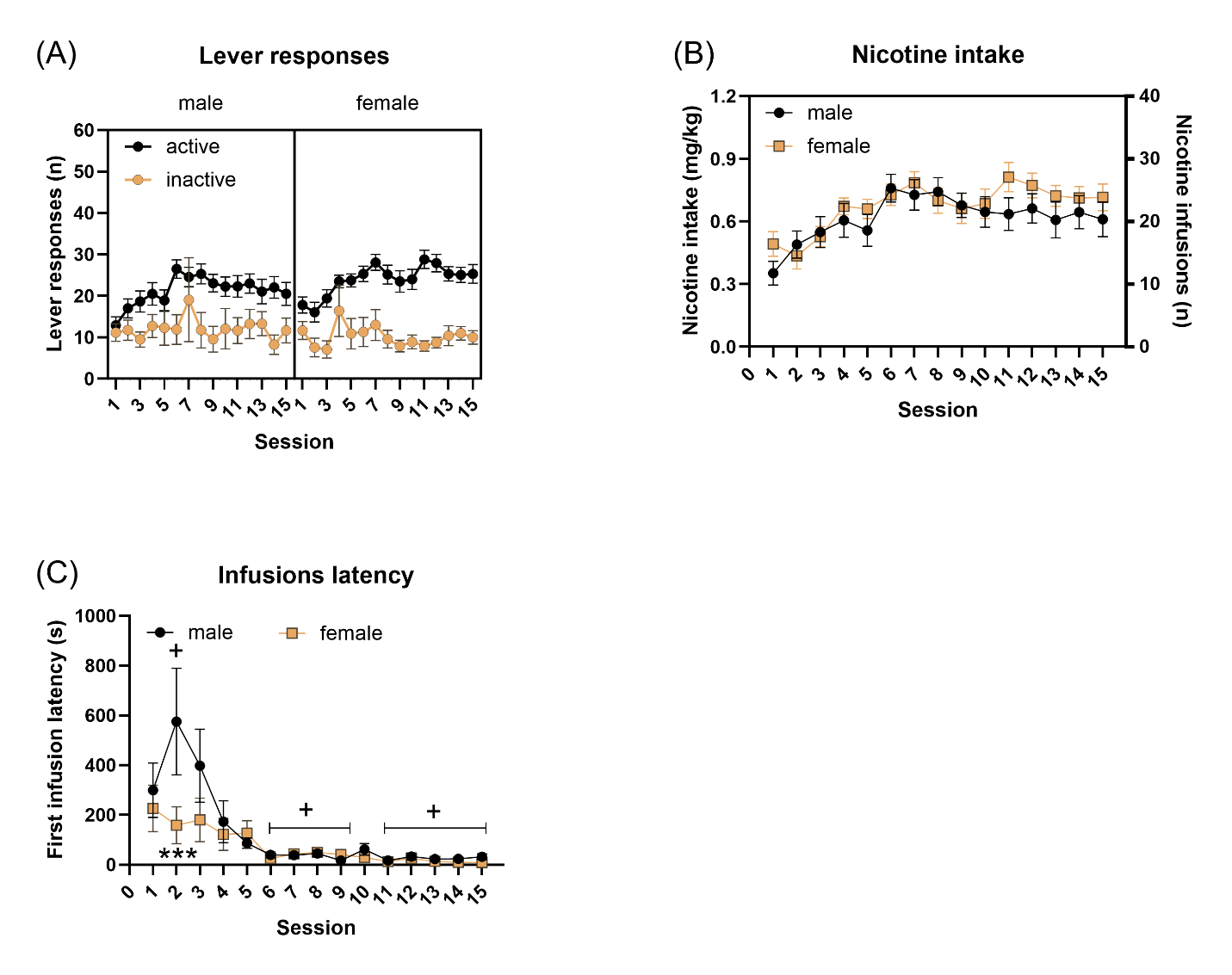

**Figure S1. Spontaneous acquisition of nicotine self-administration in male and female rats.** The rats self-administered nicotine (0.03 mg/kg/inf) during 2-hour self-administration sessions for 15 sessions. (A) Active lever responses, (B) inactive lever responses and (C) and latency to first infusion during nicotine self-administration sessions. Asterisks indicate shorter latency to the first infusion in females compared with the males on the same session day. Plus signs indicate significant difference in latency to the first infusion relative to first session in males. Males, N=12; Females, N=12. + P<0.05; *** P<0.001. Data are expressed as means ± SEM.
